# Physical and functional cell-matrix uncoupling in a developing tissue under tension

**DOI:** 10.1101/306696

**Authors:** Amsha Proag, Bruno Monier, Magali Suzanne

**Affiliations:** LBCMCP, Centre de Biologie Intégrative (CBI), Université de Toulouse, CNRS, UPS, France

## Abstract

Tissue mechanics play a crucial role in organ development. It relies on cells and extracellular matrix (ECM) mechanical properties, but also on their reciprocal interaction. The relative physical contribution of cells and ECM to morphogenesis is poorly understood. Here, we dissected the mechanics of the envelope of the *Drosophila* developing leg, an epithelium submitted to a number of mechanical stresses: first stretched, it is then torn apart and withdrawn to free the leg. During stretching, we found that mechanical tension is entirely borne by the ECM at first, then by the cellular monolayer as soon as they detach themselves from one another. Then, each envelope layer is removed by an independent mechanism: while ECM withdraws following local proteolysis, cellular monolayer withdrawal is independent of ECM degradation and driven by an autonomous myosin-II-dependent contraction. These results reveal a physical and functional cell-matrix uncoupling that could timely control tissue dynamics during development.

## INTRODUCTION

For many years, research has focused on genetic and biochemical regulation of developmental processes. More recently, the development of new approaches based on live-imaging and micromanipulation has brought novel insight into the physical properties of cells and tissues during morphogenesis, demonstrating the importance of cells and tissues mechanics during development (Monier, et al., 2015; Heisenberg and Bellaïche, 2013; Fernandez-Gonzalez and Zallen, 2011; Blankenship, et al., 2006; Bertet, et al., 2004). Tissues have viscoelastic properties that depend for the most part on the architecture and dynamics of both cytoskeletal networks and extracellular matrix (ECM) (Ingber, 2006). On the one hand, cell contractility relies mainly on the activity of acto-myosin, a macromolecular machinery composed of self-assembled actin filaments and non-muscle myosin II (Lecuit, et al., 2011). On the other hand, the extracellular matrix, which consists in a meshwork of multiple components including collagen IV, laminins and perlecan, provides support to the epithelium (Miller, 2017; Theocharis, et al., 2016).

A key question in the field is the contribution of mechanical signals during long-term development and how these signals are integrated during morphogenetic processes (Gilmour, et al., 2017; Keller, 2012). The role of mechanics in tissue deformation is beginning to be well characterized at a local scale, during relatively short periods of time and considering the epithelial sheet as a single viscoelastic entity. However, characterizing global tissue mechanics taking into account the respective role of ECM and cell layer over long timescales remains difficult due to technical limitations. In particular, how the mechanical properties of cells and the extracellular matrix integrate to confer its physical properties to the tissue remains poorly understood in living organisms (Daley and Yamada, 2013).

To characterize the respective contribution of cells and the matrix in tissue morphogenesis, we took advantage of the isolated developmental system that is the *Drosophila* leg disc. The *Drosophila* leg disc has a relatively simple organization. It is composed of two juxtaposed tissues, the peripodial envelope and the leg proper. The peripodial envelope surrounds the developing leg, and both tissues are joined together at the proximal region of the disc (Fig1a). As any mature epithelium, the envelope is composed of a cell monolayer and an underlying extracellular matrix (ECM) called the basement membrane. Furthermore, cell division is mostly absent and the pool of extracellular matrix components is not renewed (see below). Thus, with a given number of cells and a given amount of matrix, imaginal disc development constitutes a relatively simple model system to address the contribution of mechanics in a developing tissue. Interestingly, the basement membrane forms the outer most layer of the leg disc with the cell monolayer lying right underneath and constituting a very thin squamous epithelium. This configuration makes the envelope easily accessible to micromanipulation. Furthermore, imaginal discs develop normally in culture (Aldaz, et al., 2013; Aldaz, et al., 2010; Fristrom and Fristrom, 1993), indicating that they behave as independent entities whose mechanics can be characterized throughout the whole evagination process. Using this model, we dissect the mechanical properties of a tissue under tension, the leg disc envelope, and identify specific contributions and dynamics of epithelial versus matrix layers during development.

**Figure 1:**
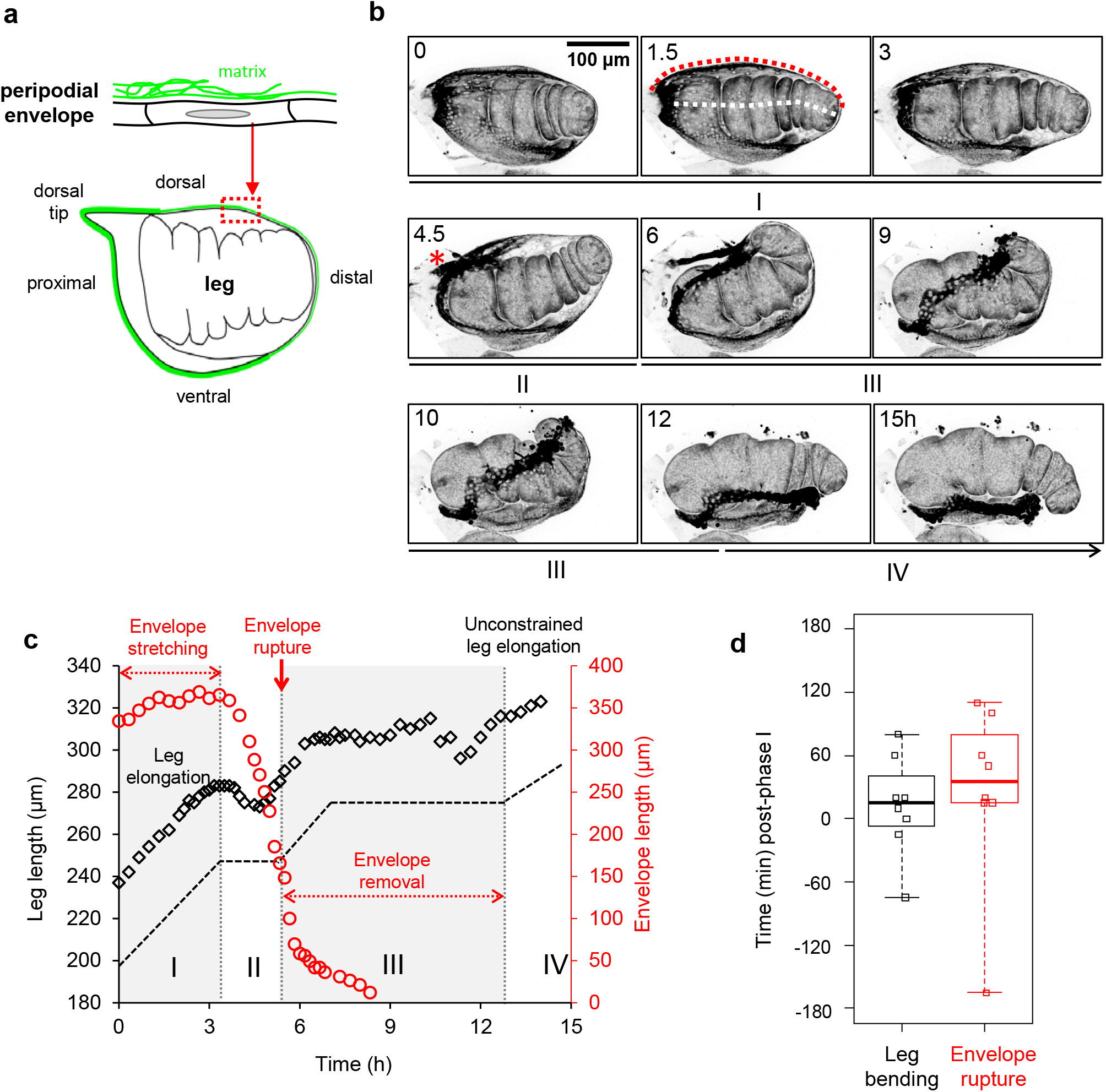
Envelope stretching, tearing and removal during leg elongation. **a.** Scheme of the leg disc at the start of elongation. The leg proper is surrounded by the peripodial envelope, composed of a thin squamous epithelial monolayer lying on a basement membrane. The basement membrane (green) is located on the outer side of the envelope. **b.** Leg disc eversion in culture. Time-lapse confocal microscopy images (z-projections) of a leg disc expressing fluorescent myosin (*sqh::sqhGFP*). A red asterisk denotes rupture at the dorsal tip. See also **Movie 1**. Representative of over 100 leg discs. Dotted red and white lines at timepoint 1h30 outline respectively the length of the envelope and of the leg measured and shown in **c.** **c.** Dynamics of leg elongation and envelope stretching. Length of the leg (black) shown in **b.** and of its envelope (red) over time. The black dashed line is a simplified view of leg elongation dynamics. Phases I-IV are described in the text. The dimensions measured on the tissue are indicated on **b.** (timepoint 1h30). **d.** Occurrence times of leg bending and envelope rupture respectively, with respect to the end of phase I. Boxes show median and quartiles and whiskers encompass all values (8 leg discs).

We show that although the matrix initially supports all the stress, the epithelium quickly detaches from the matrix and becomes highly tensed. Then, in a second phase, the envelope ruptures allowing appendage eversion. It has been proposed previously that matrix degradation by MMPs was an essential step in envelope removal in the wing disc (Srivastava, et al., 2007). However, our data reveal that the biochemical signal of MMP production, a prerequisite for subsequent eversion, is not sufficient for envelope removal. Indeed, we discover that both layers of the envelope, the matrix and the cell monolayer, rupture and withdraw independently from one another, driven by a combination of biochemical and mechanical signals: the extracellular matrix is locally degraded by matrix metalloproteases and the cell monolayer withdraws autonomously through an increase of myosin II-dependent cell contractility. Thus, these data reveal physical and functional cell-matrix uncoupling during a developmental process.

## RESULTS

### I. Envelope stretching, tearing and removal during leg elongation

During metamorphosis, leg discs evert, going from a flat to a tubular elongated structure that prefigures the adult leg (von Kalm, et al., 1995; Fristrom and Fristrom, 1993; Milner, 1977). Following the global dynamics of the envelope during leg evagination by time-lapse fluorescence microscopy, we discovered that the process is very stereotyped and proceeds through the following steps (Fig1b, FigS1 and Movie1):

– First (Fig1c, phase I), the leg elongates, a process that relies on cell shape changes and rearrangements (Condic, et al., 1991). While leg elongation progresses regularly, the peripodial envelope gradually gets more and more stretched.
– Then, leg elongation slows down to a plateau (Fig1c) and envelope stretching reaches its maximum. This stage can be divided into two different phases, before and after envelope rupture. Before rupture (phase II), the leg bends dorsally (Fig1d). Then, the envelope breaks open at the dorsal tip and starts to retract (phase III). While envelope retraction proceeds, the bending of the leg relaxes. Leg elongation remains negligible while the envelope retracts from dorsal to ventral.
– Lastly, the envelope is totally removed and leg elongation resumes (Fig1c, phase IV).

These observations strongly suggest constant mechanical interplay between the leg and the envelope. Indeed, leg elongation coincides with envelope stretching and both movements slow down at the same time, suggesting that leg elongation is responsible for envelope stretching, but that it is restrained when the envelope reaches a state of maximal tension. Then the leg bends dorsally in the direction of the future rupture site, relaxing only after the envelope breaks. Finally, once the envelope is totally removed, the leg elongates again, free from external constraint.

These observations highlight the fact that the peripodial envelope undergoes a number of mechanical stresses during leg development: it is progressively stretched, then breaks stereotypically at the dorsal tip before finally retracting. Focusing here on both the extracellular matrix and the cell monolayer of the peripodial envelope, our goal was to determine the impact of mechanics on the envelope throughout the whole process as well as the mechanical versus biochemical contribution to envelope behavior.

### II. Physical uncoupling of the ECM from the cell monolayer during envelope stretching

As mentioned above, the envelope is composed of an epithelial cell monolayer lying on ECM. Previous work has proposed that disc eversion relies on ECM degradation by matrix metalloproteases (Srivastava, et al., 2007). Yet, to date, matrix dynamics have never been followed along the whole process in a living tissue. Since the matrix contributes to tissue mechanics (Ingber, 2006), we asked whether its remodeling could alter the mechanical properties of the envelope. Thus, we analyzed the dynamics of fluorescent matrix components (collagen IV and perlecan) and observed that (1) the pool of matrix appears unchanged during leg elongation, as evidenced by the lack of fluorescence recovery several hours after photobleaching (overnight culture, not shown) and (2) the ECM is progressively remodeled during leg elongation. Initially, the matrix is rather homogeneously distributed in the whole tissue; however, as stretching progresses, the matrix becomes sparser in the distal part of the envelope. High-resolution images reveal small gaps in the matrix layer (Fig2a). This may indicate either that the concentration of matrix components was too close to background fluorescence intensity, *i.e.*, below our level of detection, or that the matrix actually ruptures in some small regions. Eventually, the matrix is locally degraded at the dorsal tip of the envelope, as shown by the appearance of a hole in the collagen IV network at the dorsal tip of the envelope layer (Fig2b and Movie 2). Strikingly, the rupture of the underlying epithelium occurs slightly later. Indeed, the hole in the matrix was always observed before epithelial rupture (Fig2b and Movie 2). This suggests that the matrix and the cell monolayer behave independently at the time of rupture. Indeed, during phase I, the ECM and the cell monolayer display correlated local deformations (Fig2c). By contrast, as from phase II, the ECM and the monolayer frequently present independent motion (Fig2d). Together, these data point towards a gradual uncoupling of the two layers from each other.

**Figure 2:**
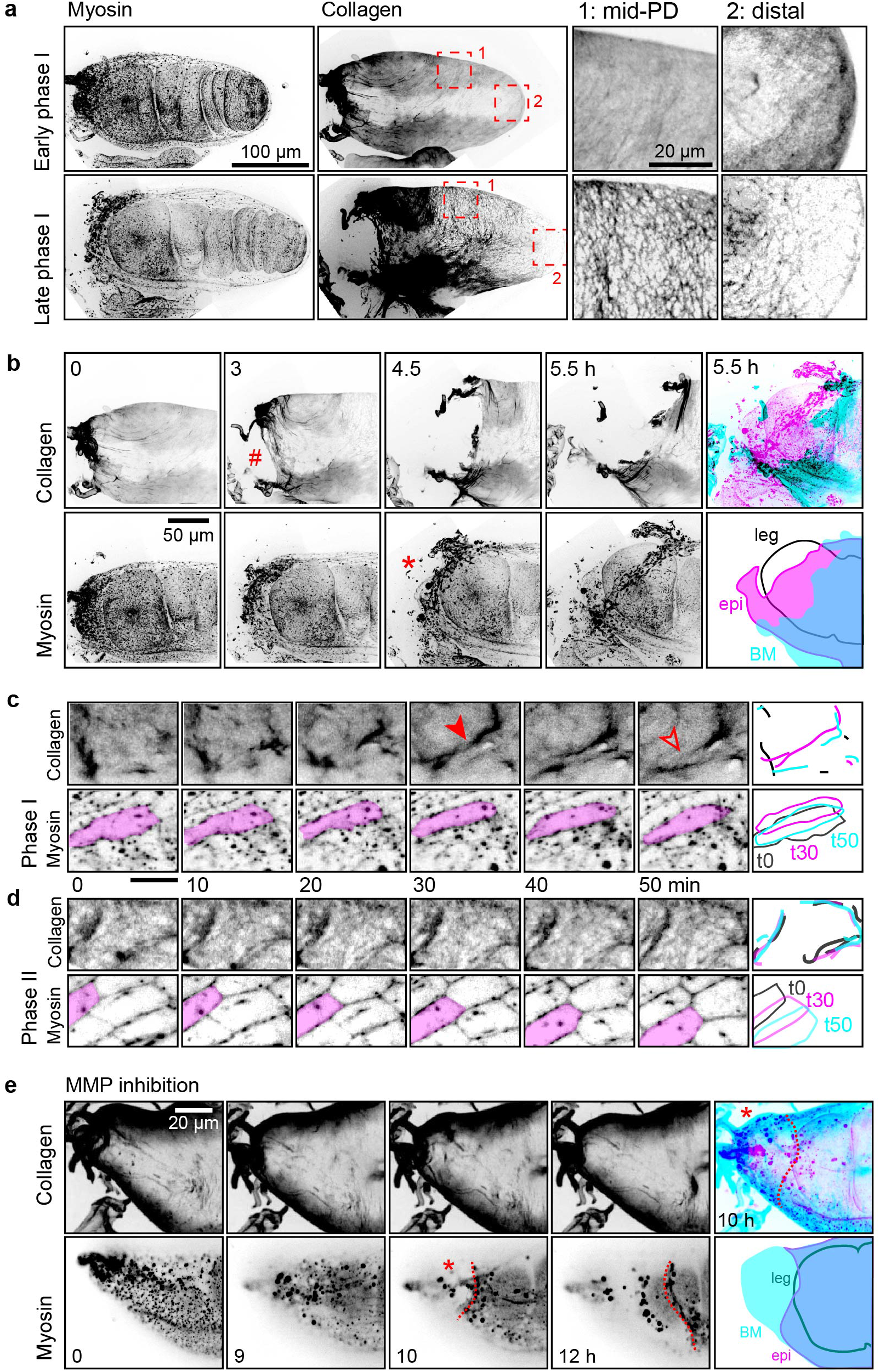
Uncoupling of the ECM from the cell monolayer during envelope stretching. **a.** Confocal images (z-projections) of a leg disc (dorsal view) expressing fluorescent collagen (*vkg-GFP*) and myosin (*sqh-TagRFPt[9B]*), at the start (upper panels) and the end (lower panels) of phase I. Right panels: enlarged views of the regions enclosed in red dashed squares, at the middle of the proximal-distal axis (1) and at the distal region (2). Note that contrast was increased in the enlarged view of region 2. Representative of 19 leg discs. **b.** Time-lapse confocal images (z-projections) of the proximal region of the same leg disc as **a.**, showing that the basement membrane ruptures before the epithelium (red hash, matrix rupture; red asterisk, epithelial rupture). Representative of 39 leg discs (collagen (*vkg-GFP*), 34, perlecan (*trol-GFP*), 5). See also **Movie 2**. Rightmost panels: merged images of collagen (cyan) and myosin (magenta) and corresponding outlines of the basement membrane (**BM**, cyan), the epithelium (**epi**, magenta) and the leg (leg, black). **c-d.** Time-lapse confocal images (single planes) of the envelope of a leg disc (dorsal view). Top and bottom rows show collagen and myosin, respectively. Rightmost panels: schematics of matrix and cell dynamics at timepoints 0, 30 and 50 min (black, magenta and cyan respectively). **c.** During phase I, the matrix displays transient wrinkle formation (filled red arrowhead, wrinkling; open arrowhead, wrinkle flattening). These dynamics are associated with cell shrinkage (compare the shape of the magenta-colored cell at timepoints 0 and 30 min). **d.** During phase II, the matrix is uncoupled from the cell layer (see magenta-colored cell), which slides over without notably deforming the matrix. Scale bars: 20 μm. **e.** Time-lapse confocal images of the proximal region of an L3 leg disc expressing fluorescent collagen (*vkg-GFP*, z-projection) and myosin (*sqh-TagRFPt[9B]*, single plane), cultured with an MMP inhibitor (GM6001, 50 μM). Although the basement membrane is not degraded, epithelial rupture at the dorsal tip still occurs (red asterisk) and the free boundary of the tissue retracts (dotted red outline). Representative of 7 leg discs. Compare with **b.**, see also **Movie 3**. Rightmost panels: merged images (z-projections) of collagen (cyan) and myosin (magenta) at the onset of epithelial rupture and corresponding outlines of the basement membrane (**BM**, cyan), the epithelium (**epi**, magenta) and the leg (**leg**, black).

### III. Independent removal of the ECM and cellular layers of the envelope

The initial hypothesis that matrix degradation by matrix metalloproteases (MMPs) may be sufficient to explain envelope rupture (Srivastava, et al., 2007) was consistent with the idea that cell-matrix attachment is essential to maintain epithelial integrity and that local matrix degradation would thus disrupt the whole envelope. However, recent work has demonstrated that a monolayer epithelium can be maintained without matrix (Harris, et al., 2012). Together with our observations, these data suggest that epithelial rupture might be regulated independently of matrix degradation. To test this idea, we prevented matrix degradation by inhibiting MMPs and followed envelope dynamics.

When MMPs are inhibited during prepupal stages, the matrix is degraded normally, progressively retracts and the leg everts (data not shown). However, when MMPs are inhibited earlier, at the end of the third larval stage (L3), local degradation of the matrix at the dorsal tip was prevented. Interestingly, in this context, although the extracellular matrix remains intact, epithelial rupture still occurs at the correct position (Fig2e and Movie 3).

These results demonstrate that matrix degradation in the envelope occurs earlier than, but is not responsible for, the rupture of the cell monolayer. This reveals that the rupture events of both layers are independent of one another, and that matrix degradation by MMPs is not required for epithelium rupture and removal. This begs the question whether an active contribution of the peripodial epithelium could be involved in this process.

### IV. Tension transfer from the ECM to the cell layer in a stretched epithelium

Since the peripodial epithelium is progressively stretched during elongation, reaching a maximal length before rupture, we reasoned that the momentary slowdown of leg elongation (Phase II in Fig1c) might indicate that the envelope is under maximum strain, preventing further elongation of the leg. Thus, rupture could be a direct consequence of tissue tension.

To test this hypothesis, we investigated the tension pattern in the peripodial epithelium at different stages of the elongation process using laser dissection. Surprisingly, during early phases we observed negligible recoil of the cytoskeleton of peripodial cells, indicating that tension in the monolayer was low (Fig3a). Note that the same laser power led to significant recoil in the cells of the leg proper, which was indicative of efficient laser cutting (not shown). This absence of tension in the peripodial epithelium was unexpected since the highly anisotropic shape of these cells from the very start of the elongation process, together with the near absence of cell division and cell intercalation, suggested that tension might already be important at this stage (McClure and Schubiger, 2005). A possible explanation would be that the duration of the process of leg elongation (3-4h *ex vivo*) might lead to cell shape stabilization in the envelope, leading to the absence of tension, even though cells appear very elongated. Indeed, cellular deformations have been shown to be stabilized by dissipation on a scale of minutes (Clément, et al., 2017).

**Figure 3:**
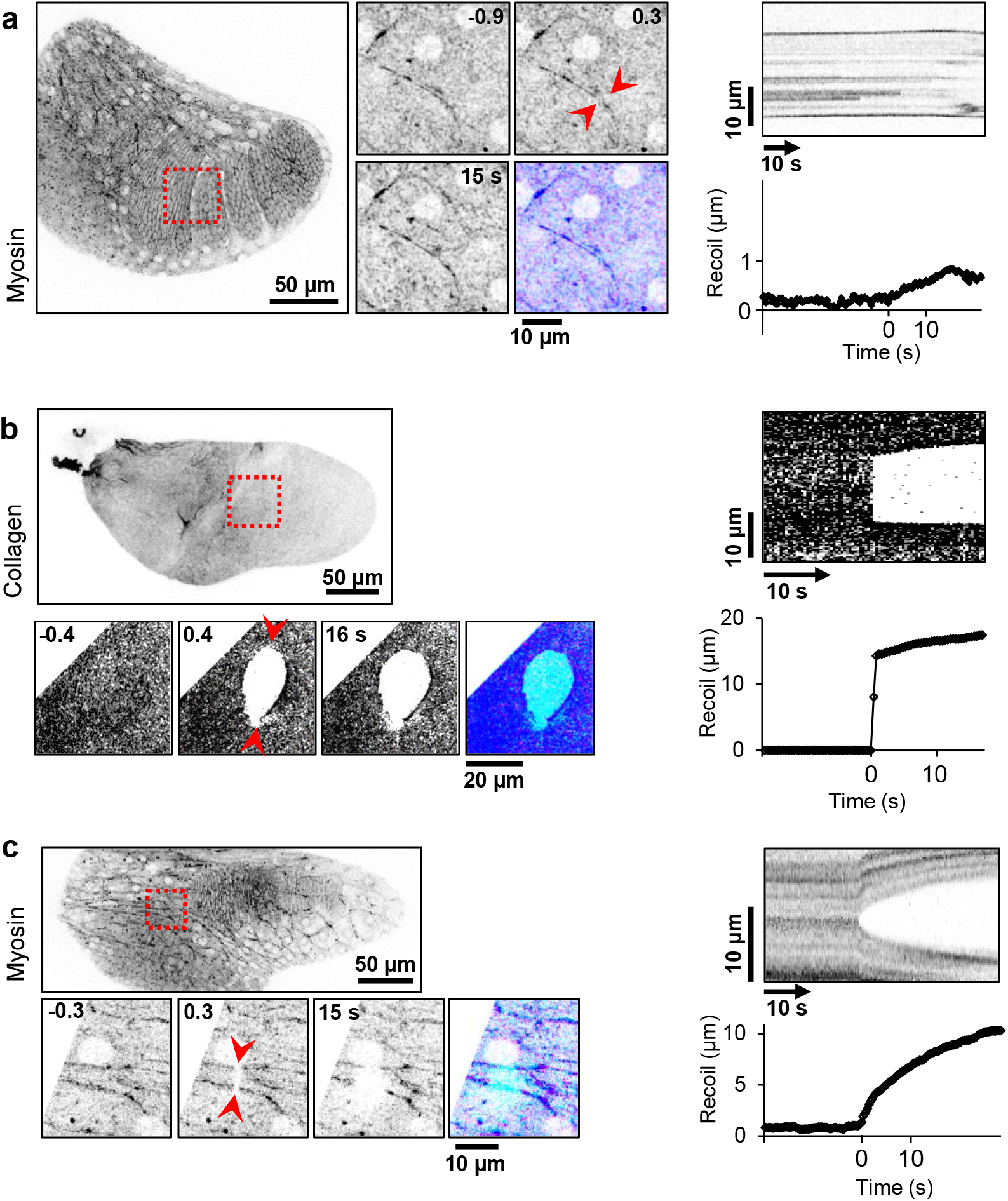
Tension transfer from the ECM to the cell layer in a stretched epithelium. **a.** The monolayer bears little tension during phase I. Left: Laser dissection of myosin filaments in cells of the envelope of a leg disc expressing fluorescent myosin (*sqh-eGFP[29B]*) in early phase I. Myosin filaments (red arrowheads) were disrupted at timepoint 0 but no recoil was observed. Bottom right image: comparison of the filament before (t = −0.9 s, cyan) and after dissection (t = 0.3 s, magenta). Representative of 5 experiments. Right: Kymograph and curve showing recoil dynamics in a representative leg disc expressing fluorescent E-cadherin. **b.** The basement membrane is under tension during phase I. Left: laser dissection of the ECM layer of the envelope of a leg disc expressing fluorescent collagen (*vkg-GFP*) in early phase I. The matrix layer was illuminated in a straight line (between the red arrowheads) at timepoint 0, leading to immediate retraction. Bottom right image: comparison before (t = −0.4 s, cyan) and at the time of dissection (t = 16 s, magenta). Representative of 5 experiments. Right: Kymograph and curve showing the recoil dynamics. **c.** The monolayer is under tension during phase II. Left: laser dissection of myosin filaments in cells of the envelope of a leg disc in phase II. The myosin network was illuminated in a straight line (between the red arrowheads) at timepoint 0 and retraction was followed over time. Bottom right image: comparison before (t = −0.3 s, cyan) and after retraction (t = 15 s, magenta). Representative of 5 experiments. Right: Kymograph and curve showing the recoil dynamics.

Alternatively, we reasoned that tension in the envelope could be entirely borne by the ECM. Given the low thickness matrix is (under 1 μm), we proceeded to cut through using high-power laser illumination to measure matrix tension. Retraction of the matrix around the cut revealed that it behaves as an elastic sheet under tension (Fig3b). These results confirmed that the envelope is under tension at the beginning of leg elongation as already shown before in other imaginal discs (Milner, et al., 1983). However, we show here that envelope tension is borne by the matrix while the underlying cell monolayer appears relaxed at this stage. Thus, the epithelial monolayer and the matrix bear different amounts of tension.

Later on, the mechanical properties of the envelope change drastically. Indeed, laser dissection revealed that tension strongly increases in the cell monolayer at the end of the elongation phase (Fig3c). Since the basement membrane is uncoupled from the cell monolayer at this stage, the monolayer must now also sustain the tension resulting from envelope stretching. This suggests that the epithelial monolayer would reach a state of maximal tension when envelope stretching is maximal at the end of leg elongation, just before rupture takes place.

### V. Active withdrawal of the cellular monolayer by myosin-dependent contraction

Interestingly, this apparent transfer of tension in the envelope coincides with a redistribution of myosin II on the dorsal side of the peripodial epithelium. While myosin II is essentially cytoplasmic at early stage, with small accumulations forming dots, it reorganizes into supracellular cable-like structures that extend radially from the future rupture site (Fig4a and Movie 4). Consistently with this redistribution of myosin II, dorsal peripodial cells contract their apex at this stage, leading to a drastic reduction of their surface and eventually of the whole dorsal region of the envelope. Importantly, the onset of envelope contraction systematically precedes its rupture (Fig4b).

**Figure 4:**
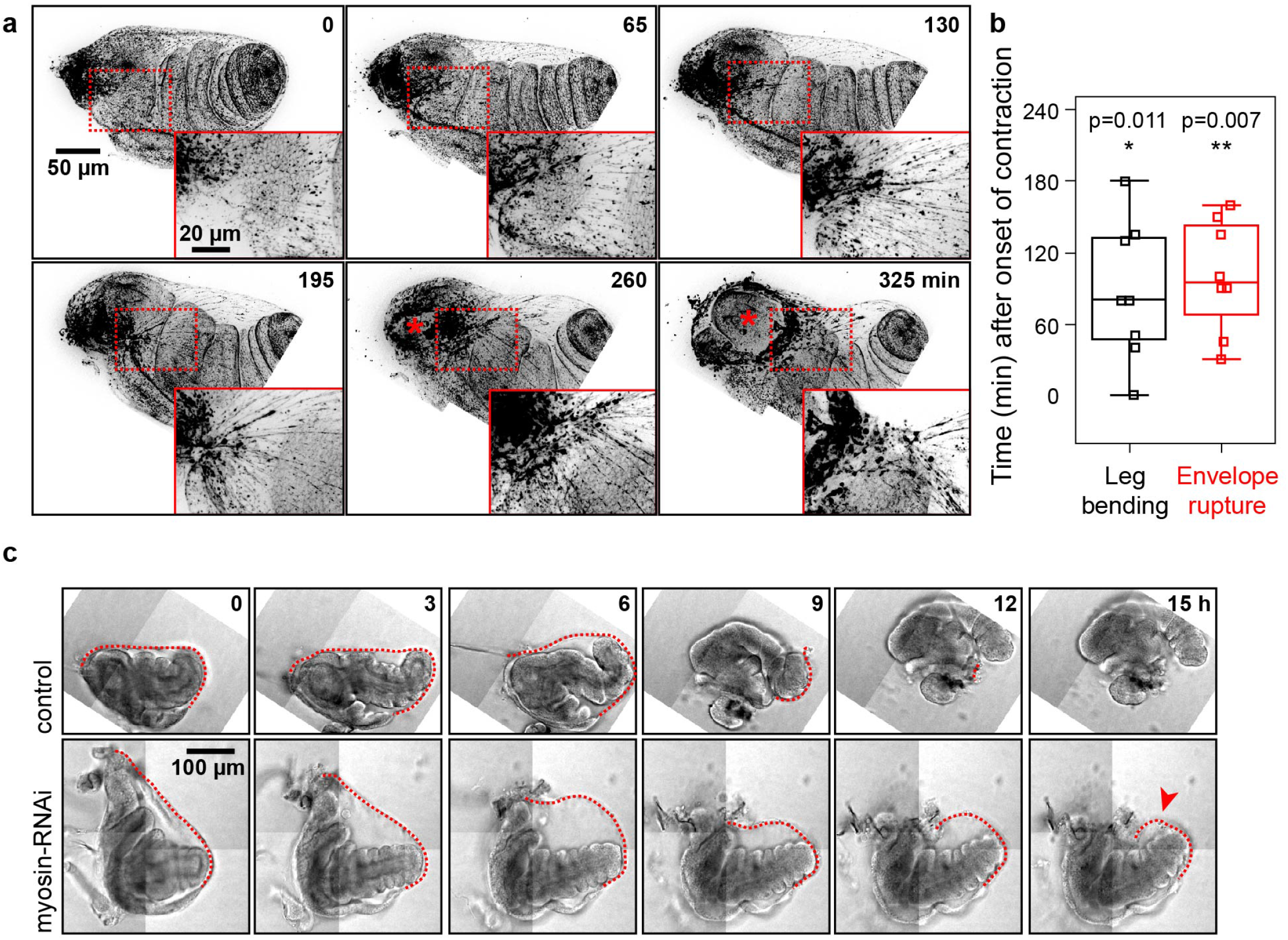
The envelope actively withdraws by myosin-dependent contraction. **a.** Time-lapse confocal images (z-projection) of the rupture site in a leg disc (dorsal view) expressing fluorescent myosin (*sqh-TagRFPt[9B]*) and showing the formation of myosin cables (red arrowheads) radially distributed just distally from the rupture site. Representative of 9 legs. Red asterisk indicates the hole in the epithelium after rupture. Insets: enlarged views of the dotted red rectangle around the myosin cables before the rupture site. Scale bar: 50 μm. See also **Movie 4**. **b.** Occurrence times of leg bending and envelope rupture respectively, with respect to the onset of envelope contraction. Boxes show median and quartiles and whiskers encompass all values (8 leg discs). p-values indicate the statistical significance that these events occur after contraction, as assessed by the one-sided Wilcoxon signed-rank test between the absolute time values paired per leg disc. **c.** Time-lapse bright-field images of a leg disc expressing myosin-RNAi in the envelope (*c855a-Gal4 > UAS-zip-RNAi*, bottom row) compared to a control leg disc (*UAS-zip-RNAi*, top row). Myosin depletion prevents retraction of the peripodial envelope (red arrowhead). Representative of 7 legs. Red dashed line outlines the envelope over time.

Altogether, these results indicate that the mechanics of the peripodial epithelium change drastically during leg elongation, in terms of both tension and contractility. Indeed, at the time of rupture, dorsal peripodial cells appear to be uncoupled from the matrix, under high tension and strongly contractile.

Since cell contraction in the envelope occurred just prior to epithelial rupture, we asked whether contraction could be involved in epithelial rupture and/or removal. To test this hypothesis, we attempted to inhibit contraction by targeting myosin II. Strongly inhibiting myosin II in the peripodial epithelium by genetic means (expressing dominant negative forms or RNAi) tended to alter epithelial integrity. Therefore, we set up conditions in which contraction was visibly affected without notably perturbing the general organization of the tissue. In these conditions, we observed that although rupture appears to take place normally, envelope removal is prevented (Fig4c). This experiment suggests that contraction is specifically required for envelope removal.

## DISCUSSION

In this study, we analyzed the behavior of an epithelium under tension, revealing important physical dynamics. We focused on the envelope of the *Drosophila* imaginal leg disc, which undergoes a series of modifications during leg development, including stretching, rupture and retraction.

During envelope stretching, we observed that tension in the envelope is first borne by the matrix, then transferred to the epithelial layer as it loses its interaction with the extracellular matrix. During this second step, peripodial epithelium breaks open independently of matrix degradation. These results bring to light a dynamical process of developmental epithelial tearing, supporting a model in which cell-matrix disengagement leads to matrix-to-cell tension transfer in order to trigger an active retraction of the cell monolayer (Fig5). Thus cell-matrix uncoupling could act as a developmental timer, and hence constitute an alternative to classical hormonal signals for the control of stereotyped organ morphogenesis.

**Figure 5:**
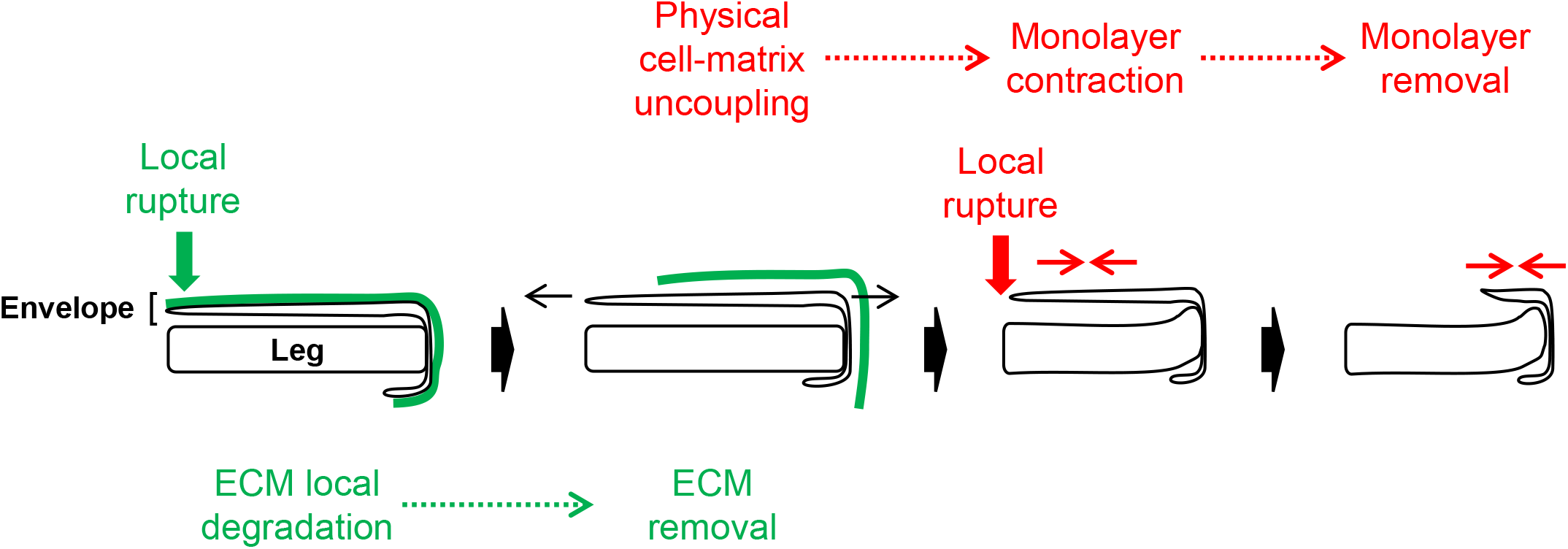
Model of the biochemical and physical processes involved in leg disc envelope removal. Both layers of the envelope behave independently in terms of tension, rupture and withdrawal. The ECM (green layer) undergoes local proteolysis early on, before rupturing and physically detaching from the cell monolayer (green sequence). The monolayer then increases its contractility, ruptures and then retracts through autonomous contraction (red sequence).

Overall, these results reveal that epithelia do not necessarily lose their integrity when they lose matrix adhesion. Indeed, at least under tension, they can conserve their structure and survive independently, as previously shown for cultured cells (Wyatt, et al., 2015; Harris, et al., 2012). The influence of spreading area on cell proliferation and apoptosis has been studied for decades now (Chen, et al., 1997) and it is clear that in single adherent cells, low spreading area favors cell death. However, in a monolayer epithelium, it is not known whether the maintenance of intercellular adhesion might compensate for basal adhesion loss, hence promoting cell survival through a combination of signaling from adherens junction-linked proteins and mechanotransduction (Discher, et al., 2009; Janmey, et al., 2009). Further studies are needed to determine if this is a general feature of epithelia under tension.

Most studies on the mechanical properties of epithelia to date have focused either on the epithelial cells or on the matrix. However, both are essential constituents of epithelia, differing notably in their composition and mechanical response and conferring particular physical properties to tissues and organs, which highlights the importance of characterizing both. Each cell can modify its shape and rigidity actively through the reorganization of cytoskeletal components and the generation of intracellular forces by molecular motors such as myosin. As for the matrix, since it does not contain any motor proteins, it is considered as a passive element, its rigidity depending directly on its composition and architecture. Here, we benefitted from the particular geometry of the envelope (a flat epithelium supported by a basement membrane forming the outer layer of the disc and thus directly accessible) and from the fact that leg tissues can accomplish their morphogenesis in culture. Thus, we could decipher the relative contribution of matrix and epithelial cells to tissue mechanics during a long-scale process of morphogenesis. Combining live imaging with biophysical approaches, we discovered that both layers of the envelope behave independently in terms of tension, rupture and withdrawal, responding to different signals, either biochemical or mechanical. Thus, under high tension, epithelial cells and their basement membrane appear to uncouple their responses, most probably due to the difference between their physical properties. Surprisingly, these results reveal that the layers constituting an epithelium, although generally viewed as interdependent, can behave independently under certain circumstances, highlighting the necessity to revisit the classical vision of epithelia.

## Material and Methods

### Fly stocks

sqh-eGFP[29B] and sqh-TagRFPt[9B] (this work) are knock-in designed and generated by homologous recombination by InDroso functional genomics (Rennes, France). The respective tags were inserted in C-terminal just before the stop codon and the resulting flies were validated by sequencing. sqh::sqh-GFP on the 2nd chromosome (Royou, et al., 2002) was already described. vkg-GFP[G0454] and Trol-GFP[G00022] are Flytrap lines (Morin, et al., 2001). C855a-Gal4 and uas::zip-RNAi[7819] were obtained respectively from BDSC and VDRC.

### Sample preparation

Leg discs were dissected from L3 larvae, white pupae or 2h APF prepupae in Schneider’s insect medium (Sigma-Aldrich) supplemented with 15 % fetal calf serum and 0.5 % penicillin-streptomycin as well as 20-hydroxyecdysone at 2 μg/mL (Sigma-Aldrich, H5142). Leg discs were transferred on a glass slide in 13.5 μL of this medium confined in a 120 μm-deep double-sided adhesive spacer (Secure-Seal^TM^ from Sigma-Aldrich) and a glass coverslip was then placed on top of the spacer. A precision glass coverslip (from Marienfeld, Germany) was used for laser dissection experiments. Halocarbon oil was added on the sides of the spacer to prevent dehydration. Dissection tools were cleaned with ethanol before dissection. For MMP inhibition experiments, leg discs were mounted in medium supplemented with GM6001 (gift from E. Théveneau, CBD, Toulouse, France) (50 μM, 0.5 % DMSO). For myosin inhibition experiments, flies were crossed at 25°C and larvae were grown at 18°C to limit Gal4 activity and avoid disrupting tissue integrity.

### Live microscopy and imaging

Leg disc development was imaged using an inverted spinning disk confocal microscope (CSU-X1, Yokogawa, coupled to a Leica (resp. Zeiss) microscope) mounted with 20x/0.8 multi-immersion or 40x/1.2 oil objectives (resp. 20x/0.8 air objective) and equipped with 488 nm and 561 nm LEDs and a piezo stage. Images were acquired over time with a Hamamatsu EMCCD camera controlled by the Metamorph (resp. Zen) software, at a rate of one z-stack every 5 to 15 min. Images were processed with the ImageJ software for registration (StackReg plugin), bleaching correction (Bleach Correction plugin), background correction and smoothing. The length of the envelope and the leg (Fig1c and FigS1) was measured on image z-stacks, from the dorsal tip to the distal pole (envelope) and from the femur to the distal pole (leg) as indicated on Fig1b. The occurrence time of a particular event (Fig1d and Fig4b) was defined as the first timepoint when the event was visible.

### Laser dissection

Laser dissection experiments were performed with a pulsed DPSS laser (532 nm, pulse length 1.5 ns, repetition rate up to 1 kHz, 3.5 μJ/pulse) steered by a galvanometer-based laser scanning device (DPSS-532 and UGA-42, from Rapp OptoElectronic, Hamburg, Germany). The laser beam was focused through an oil-immersion lens of high numerical aperture (Plan-Apochromat 63x/1.4 Imm Oil or LD LCI Plan-Apochromat 63x/1.2 multi-Imm, from Zeiss). Photo-disruption was produced in the focal plane by illuminating at 60 % laser power. To target intracellular structures, illumination duration was set at 40 ms on a small spot focused on a filament or junction at zoom 2x. For the matrix layer, 20-40 μm-long lines were illuminated for 50-100 ms at zoom 1x. Images were acquired every 200 ms to 400 ms during the experiment using a confocal laser scanning microscope (LSM-880, Zeiss) equipped with a 488 nm Argon laser and a GaAsP photomultiplier. Data analysis was performed with the ImageJ software using a home-made macro.

### Statistics

The statistical significance of the difference between occurrence times of particular events (Fig4b) was assessed using the one-sided Wilcoxon signed-rank test. Absolute time measurements were paired by leg disc. The null hypothesis was that the time difference values between envelope rupture (resp. leg bending) and envelope contraction were samples from a symmetric distribution centred below 0. Calculations were performed with R and p-values are indicated on the graph.

## Author contribution

MS, BM and AP designed experiments. AP performed experiments and data analysis. BM constructed fly stocks. MS supervised the project.

## Acknowledgements

The authors express their thanks to E. Théveneau for MMP inhibitors, to T. Mangeat for help with laser dissection and to all team members for helpful discussion. They are grateful to J. Fouchard and G. Charras for insightful discussions.

MS’s lab is supported by grants from the European Research Council (ERC) under the European Union Horizon 2020 research and innovation program (grant number EPAF: 648001), the Fondation Arc pour la Recherche sur le Cancer (CA 09-12-2014) and from the Institut National de la Santé et de la Recherche Médicale (Inserm, Plan cancer 2014-2019).

## Supplementary figure legends

**Figure S1:**
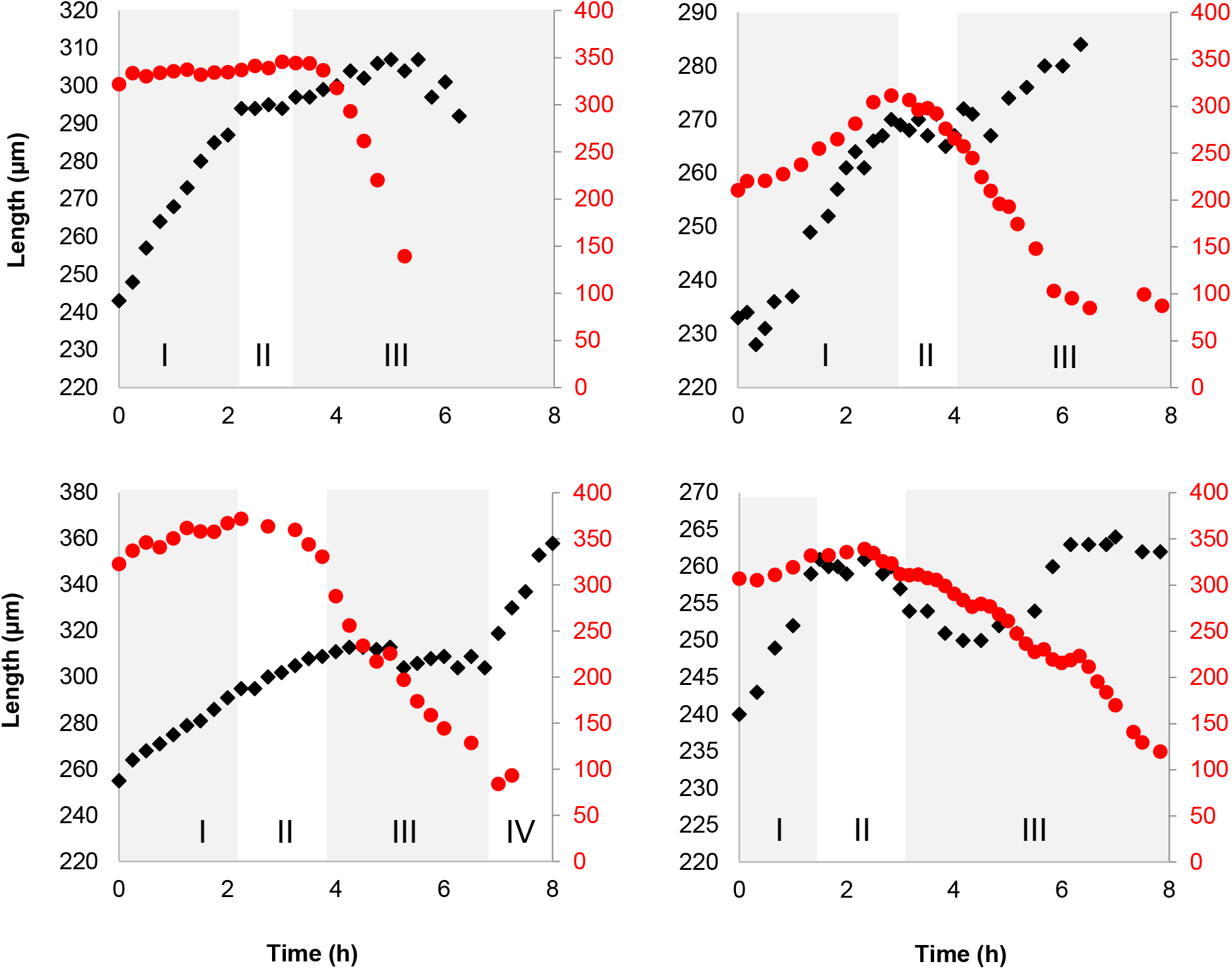
Related to Figure 1. Envelope rupture and withdrawal are coordinated with leg disc elongation dynamics. Dynamics of leg elongation and envelope stretching for 4 leg discs. Length of the leg (black) and of the envelope (red) over time, measured as indicated on Fig1b. Phases I-IV are described in the text.

**Movie 1. Related to Figure 1b. Envelope stretching, tearing and removal during leg elongation**. Leg disc eversion in culture. Time-lapse confocal microscopy images (z-projections) of a leg disc expressing fluorescent myosin. Tissue orientation follows Fig1a (dorsal-ventral from top to bottom, proximal-distal from left to right). Image stacks were acquired every 10 min.

**Movie 2. Related to Figure 2b. Uncoupling of the ECM from the cell monolayer during envelope stretching**. Time-lapse confocal images (z-projections) of a leg disc expressing fluorescent collagen and myosin during phase I. Image stacks were acquired every 7.5 min.

**Movie 3. Related to Figure 2e. Matrix degradation is not required for envelope rupture**. Time-lapse confocal images (z-projections) of the proximal region of an L3 leg disc expressing fluorescent collagen and myosin, cultured with an MMP inhibitor. Image stacks were acquired every 15 min.

**Movie 4. Related to Figure 4a. Myosin in the envelope organizes into cable-like structures before contraction and rupture**. Time-lapse confocal images (z-projection) of the rupture site of a leg disc expressing fluorescent myosin. Image stacks were acquired every 5 min.

